# Cell-generated mechanical forces play a role in epileptogenesis after injury

**DOI:** 10.1101/2025.02.09.637325

**Authors:** Laya Dalir, Svetlana Tatic-Lucic, Yevgeny Berdichevsky

## Abstract

Traumatic brain injury (TBI) is associated with a significantly increased risk of epilepsy. One of the consequences of severe TBI is progressive brain atrophy, which is frequently characterized by organized tissue retraction. Retraction is an active process synchronized by mechanical interactions between surviving cells. This results in unbalanced mechanical forces acting on surviving neurons, potentially activating mechanotransduction and leading to hyperexcitability. This novel mechanism of epileptogenesis was examined in organotypic hippocampal cultures, which develop spontaneous seizure-like activity in vitro. Cell-generated forces in this model resulted in contraction of hippocampal tissue. Artificial imbalances in mechanical forces were introduced by placing cultured slices on surfaces with adhesive and non-adhesive regions. This modeled disbalance in mechanical forces that may occur in the brain after trauma. Portions of the slices that were not stabilized by substrate adhesion underwent increased contraction and compaction, revealing the presence of cell-generated forces capable of shaping tissue geometry. Changes in tissue geometry were followed by excitability changes that were specific to hippocampal sub-region and orientation of contractile forces relative to pyramidal cell apical-basal axis. Results of this study suggest that imbalanced cell-generated forces contribute to development of epilepsy, and that force imbalance may represent a novel mechanism of epileptogenesis after trauma.

## Introduction

Traumatic brain injury (TBI) is associated with a significantly increased risk of epilepsy (Annegers et al., 1998; Karlander et al., 2021; Pease et al., 2022). Post-injury imaging in patients with severe TBI reveals progressive brain atrophy and ventricle enlargement (Bergeson et al., 2004; Bigler and Maxwell, 2011), while imaging of animal models of TBI demonstrates progressive increase in the volume of the cavity under the impact site (Bramlett and Dietrich, 2002; Dixon et al., 1999; Pierce et al., 1998). This is accompanied by gradual retraction of neighboring brain regions containing surviving neurons (Hall et al., 2005; Thompson et al., 2005). Development of posttraumatic epilepsy occurs on a similar time scale as brain tissue atrophy and retraction. Hippocampus is particularly vulnerable to injury after TBI (Bigler et al., 1997; Maxwell et al., 2003; Wilde et al., 2007), and hippocampal atrophy is associated with posttraumatic epileptic focus in the temporal lobe (Gupta et al., 2014; Hudak et al., 2004). In animal models of TBI, regions of the brain neighboring the lesion are characterized by hyperexcitability and may contribute to generation of spontaneous seizures (Bragin et al., 2016; D’Ambrosio et al., 2004). These correlations between temporal and spatial features of injury-related atrophy and the development of epilepsy suggest a potential link between these processes. Several mechanisms behind this link, such as injury-related inflammation and network reorganization, have been proposed (Hunt et al., 2013; Pease et al., 2024). Mechanical force imbalance in the atrophying tissue may also play a role, and represent a novel mechanism of epileptogenesis after injury. While atrophy is driven by several cell death mechanisms in response to injury (Bramlett and Dietrich, 2015), the retraction of tissue after atrophy may also be a result of active cellular processes, but in surviving cells. Brain tissue retraction occurs in a seemingly organized fashion – the edge of retracting tissue is well defined, and moves gradually with time. This suggests that retraction is a synchronized behavior of the surviving cell population, resembling an inversion of the tissue expansion that occurs during development due to cell proliferation. Self-organization of healthy tissue is coordinated in part through mechanical mechanisms: tension and contraction of individual cells’ cytoskeleton is transmitted to neighboring cells through cell-cell and cell-extracellular matrix (ECM) contacts, mediated by cadherin and integrin receptors (Laurent et al., 2017). These mechanisms contribute to tissue mechanical properties (Ayad et al., 2019), result in sorting of different types of cells (Athanasiou et al., 2013), and play a role in generation of cortical folds (Llinares-Benadero and Borrell, 2019). In the developed brain, a mechanical force balance may exist between tension due to forces generated by cytoskeletons of neuronal and glial cells, and hydrostatic pressure (Essen, 1997). A similar, internally generated force balance, termed mechanical or tensile homeostasis, exists in many other tissues (Ingber, 2003; Stamenović and L. Smith, 2020). The process of tissue retraction in the brain may be a response to tissue-scale imbalances in cell-generated forces that are caused by cell death after injury. Dissociated cortical neurons and glial cells readily form aggregates which undergo contraction and compaction, demonstrating that these cells are capable of generating tissue-scale mechanical forces (Hasan and Berdichevsky, 2021). In addition to generating mechanical forces, neurons also respond to externally applied forces via a process termed mechanotransduction. Neurons generate intracellular calcium transients when mechanically stimulated (Duncan et al., 2021; Gaub et al., 2020), and are sensitive to localized forces in the low pico Newton range (Falleroni et al., 2022). Neuronal mechanotransduction occurs via several different mechanisms, which include sensation transmitted through cytoskeleton tension, mechanosensitive ion channels, or altered transcription in mechanically deformed nucleus (Pillai and Franze, 2024). Transduction of mechanical stimuli leads to changes in neuronal firing, suggesting altered excitability (Kasuba et al., 2024). Regions of neuronal aggregates that have experienced the most contraction are characterized by highest excitability, demonstrating a link between cell-generated forces and neural activity (Hasan and Berdichevsky, 2021). Taken together, these findings suggest that after traumatic brain injury, acute and secondary cell death result in mechanical imbalances that lead to compaction of surviving cells and retraction of the surviving tissue. This in turn mechanically stimulates surviving neurons, leading to hyperexcitability.

In this work, we examine whether ex vivo hippocampal tissue, with organized neurons and glial cells and native extracellular matrix, experiences cell-generated mechanical forces that shape the tissue and lead to excitability changes. We used organotypic hippocampal cultures for this study. Cultured hippocampus slices survive for 4 or more weeks in vitro. Slices experience an initial wave of cell death in the slice that lasts approximately 3 days after dissection (Berdichevsky et al., 2012). This is followed by appearance of spontaneous seizure-like activity approximately 7-10 days after dissection (Berdichevsky et al., 2013), and organotypic hippocampal cultures are used as an in vitro model of epileptogenesis (Lau et al., 2022; Liu et al., 2019). We have previously observed that organotypic hippocampal cultures thinned and underwent reduction in area in the first week in vitro. This process is likely driven by two linked mechanisms: (1) clearance of dead cells from the slice by microglia (Balena et al., 2023), and (2) imbalance in the mechanical forces between surviving cells, causing tissue contraction and retraction of tissue edges. Here, we examine whether tissue contraction is driven by cellular forces, whether introduction of further imbalances in cellular forces leads to more tissue contraction, and whether these processes alter neuronal excitability.

## Results

### Natural contraction of organotypic slices

We cultured organotypic hippocampal slices on entirely PDL-coated culture dishes to observe the natural morphological changes in the slice over time. Slice area increased from day in vitro (DIV) 0 to 1, as the slice attached to the adhesive culture substrate (Supplementary Fig. 1). Bright-field images captured on days in vitro (DIV) 1, 3, and 7 reveal further dynamic changes in the size of the slice (Fig. 1A). Tracking of the boundary of the slice (Fig. 1B) shows that significant reduction of the hippocampal culture area occurred between DIV 1 and 7, and that area stabilized between DIV 7 and 18 (Fig. 1C). These results are consistent with the presence of contractile forces in hippocampal cells.

**Fig. 1.**
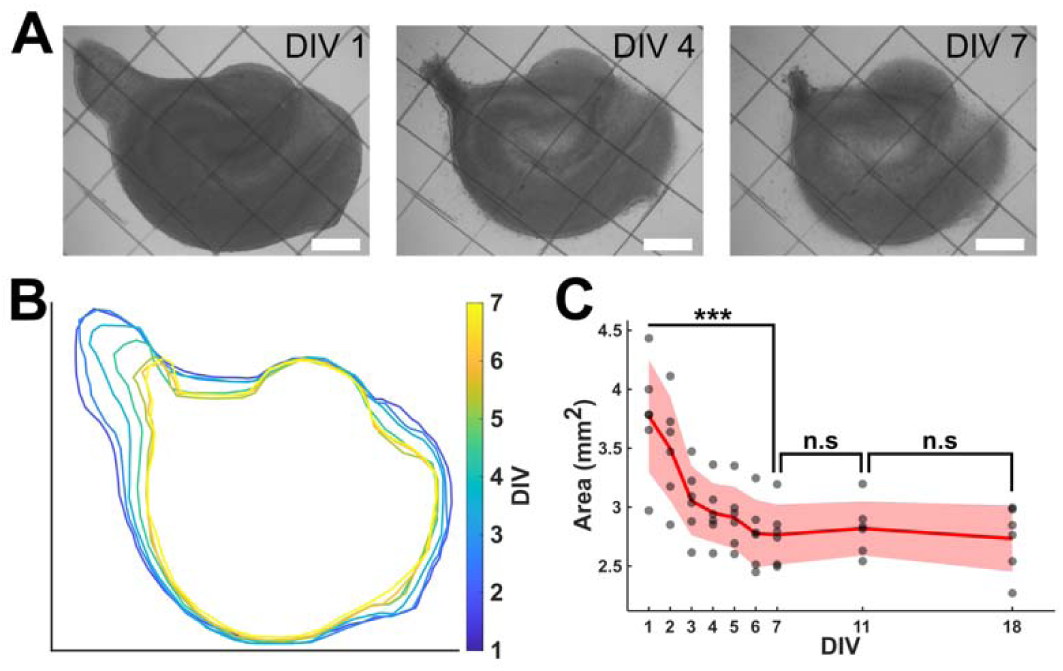
Natural contraction of organotypic slices. (A) Bright field image of cultured slices on PDL-coated surface from DIV 1 to DIV 7 (scale bar 500 µm). (B) Boundary of slice from DIV 1 to DIV 7. (C) Calculated area of cultured slices (*n* = 6) from DIV 1 to DIV 18. The red solid line and shaded area highlight the mean and standard deviation, respectively. Paired t-test was used to measure statistical significance (*** p < 0.001, n.s: not significant).

### Contractile forces are present in hippocampal tissue

Forces exerted by cells can be visualized by traction force microscopy (Polacheck and Chen, 2016). In this technique, cells are placed in an ECM gel with embedded beads. Cellular forces are transmitted to the ECM, and cause movement of beads, from which direction and magnitude of cellular forces can be estimated. To demonstrate that hippocampal cells are capable of exerting contractile forces, hippocampal slices were embedded in a mixture of Matrigel and fluorescent microbeads (Fig. 2A). Microbeads were tracked for two days after the addition of Matrigel. Bead trajectories show predominant movement toward the slice confirming the presence of tissue-driven matrix pulling (Fig. 2C). The movement angle of each bead was measured relative to the line connecting its initial location to the center of the slice (Fig. 2B). Polar histograms reveal that distribution of beads movement angle was centered around ∠ 0° (Fig. 2D, E). These results show that cells in the slice exerted traction forces on the surrounding gel. Movement of beads toward the slice demonstrates that the forces were contractile.

**Fig. 2.**
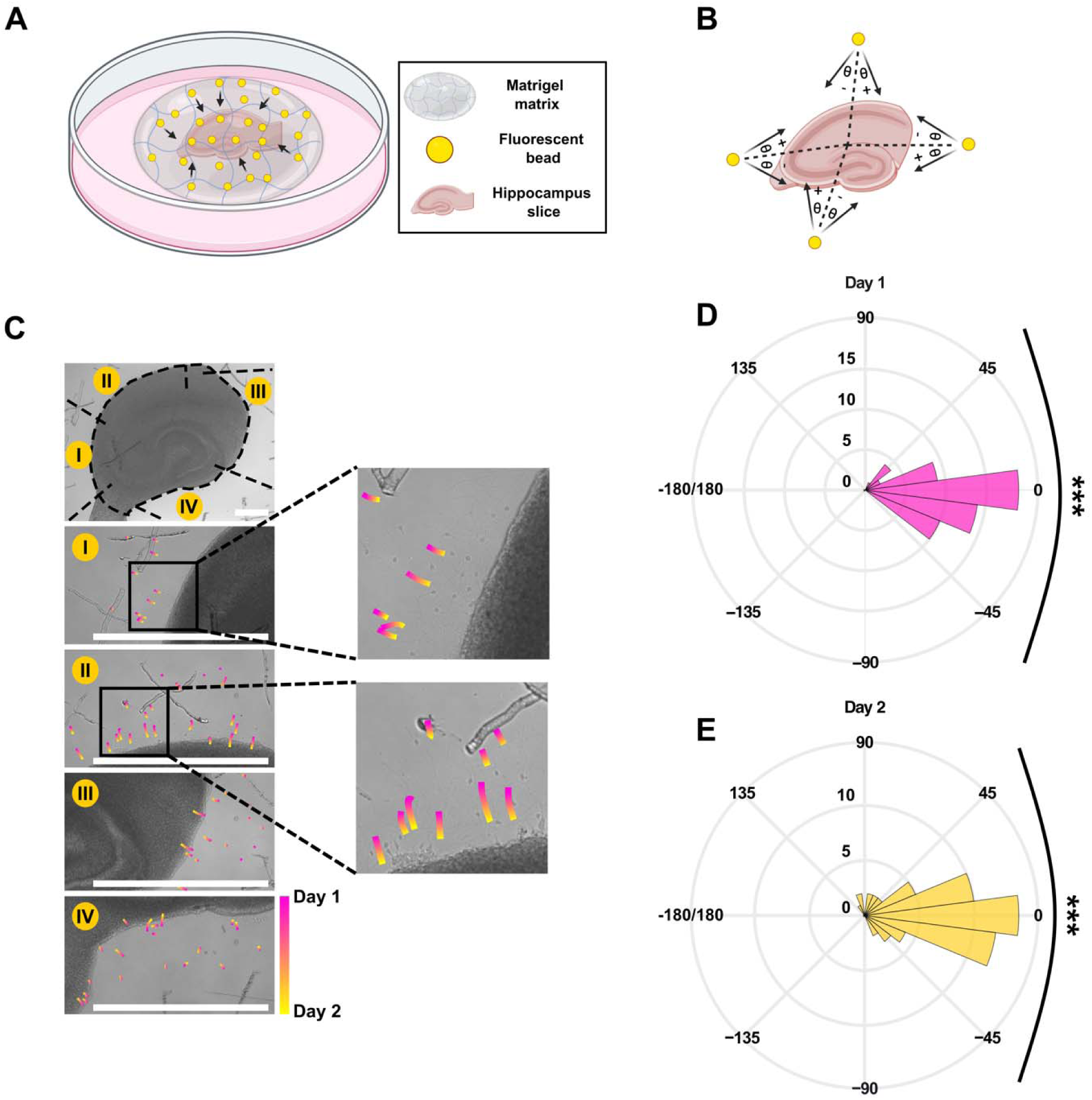
Tissue-induced matrix pulling. (A) Schematic of the organotypic slice co-cultured with Matrigel and beads mixture. (B) Beads movement angle measurement method. The angle of movement is determined as the angle formed between the bead trajectory for each day and the imaginary line represented by the black dashed line connecting the bead’s initial position to the center of the slice. (C) The top panel displays a 4X bright field image of the slice, while panels I-IV depict corresponding sections in the top panel at 10X magnification with beads trajectories. Pink color indicates movement from DIV 0 to DIV 1 and yellow from DIV 1 to DIV 2. All scale bars in this figure resemble 500 μm. (D), (E) Polar histogram of beads movement angle on day 1 and 2, respectively. The chi-squared goodness-of-fit test was conducted using 15° bins against a uniform distribution (*n* = 59, *p* < 0.001).

### Accelerated localized contraction in the slice

We varied the location and extent of hippocampal slice contraction by selectively eliminating tissue-substrate adhesion. Polydimethylsiloxane (PDMS) film was used for patterning the culture substrate coating with the cell-substrate adhesion molecule poly-D-lysine (PDL) (Fig. 3A). Different slice configurations were tested on the PDL-free areas to enhance contraction in various directions within the slice (Fig. 3B-D). We accelerated contraction in intact CA3 along the apical-basal axis (axial) (Fig. 3B), in lesioned CA3 along and transverse to the apical-basal axis (axial and transverse) (Fig. 3C), and in subiculum (SUB) transverse to the apical-basal axis (transverse) (Fig. 3D). Contraction did not affect the neuronal viability as density of CA3 neurons was not significantly different in control and contracted groups (Fig 4A-D). Control slices were cultured on entirely PDL-coated dishes. Significant increase in contraction was observed in the contracted group compared to the control group across all targets: 52% in intact CA3 (Fig. 3E), 65% in lesioned CA3 (Fig. 3F), and 69% in SUB (Fig. 3G).

**Fig. 3.**
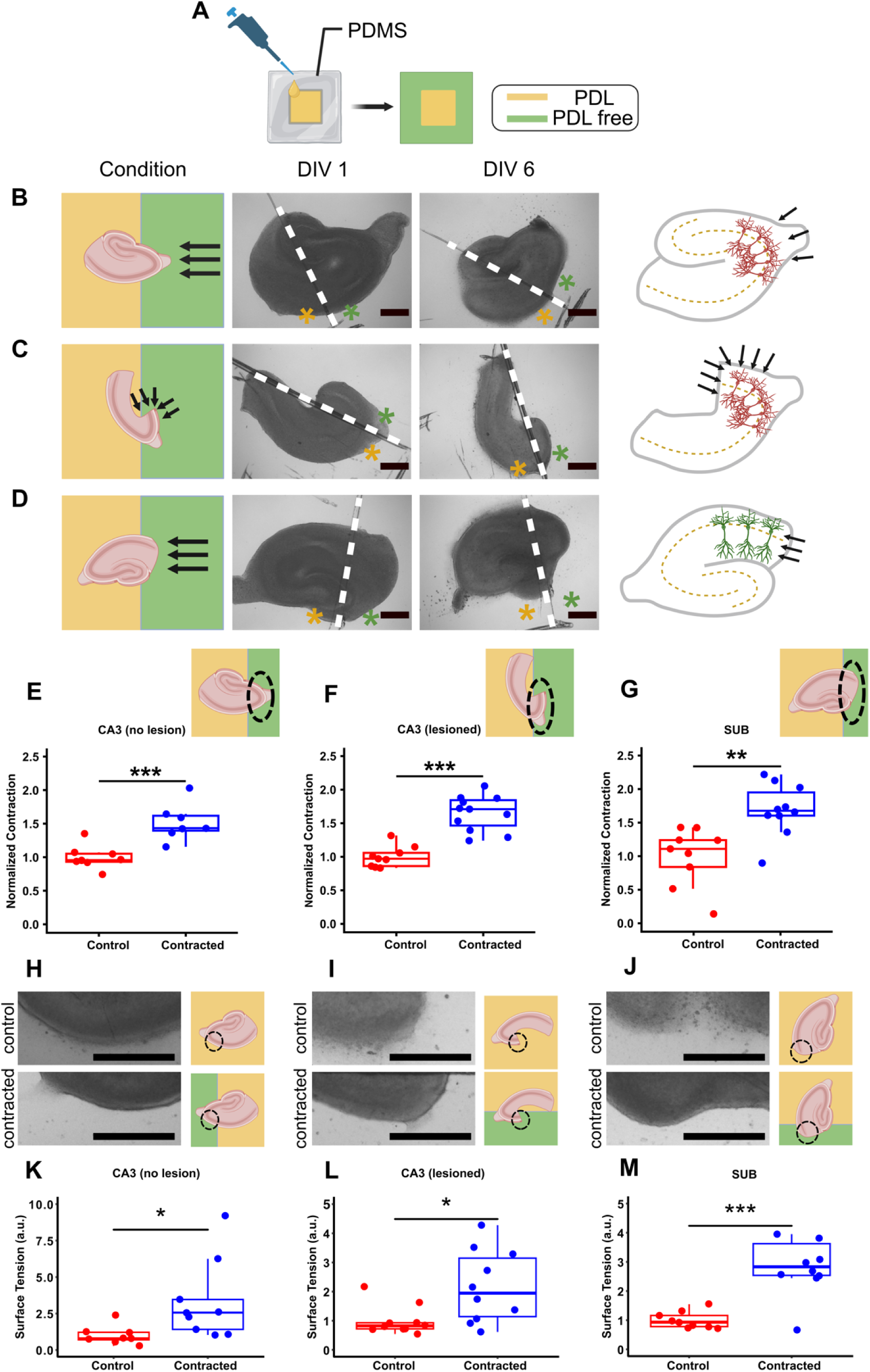
Accelerated localized contraction in the slice. (A) Schematic of the PDL-coating step. In this figure, the light-yellow color represents the PDL-coated surface, while the light green color represents the PDL-free portion of cultrure substrate. Black arrows in B, C, and D indicate contractile force direction. (B) from left to right: configuration of slice with intact CA3 on PDL-free portion of the substrate, bright field image of slice on DIV 1, bright field image of slice on DIV 6, and force direction relative to pyramidal neurons (axial). Green and yellow asterisks in the bright-field images represent PDL-free and PDL-coated portions of the substrate, respectively. (C) From left to right: configuration of slice with lesioned CA3 on PDL-free portion of the substrate, bright field image of slice on DIV 1, bright field image of slice on DIV 6, axial and transverse force direction. (D) From left to right: configuration of slice with subiculum on PDL-free portion of the substrate, bright field image of slice on DIV 1, bright field image of slice on DIV 6, and transverse force direction. All scale bars in this figure indicate 500 µm. (E), (F), (G) Normalized contraction of the slice in three different configurations: intact CA3 (*n* = 8 control, *n* = 7 contracted), lesioned CA3 (*n* = 9 control, *n* = 11 contracted), and SUB (*n* = 9 control, *n* = 10 contracted) on PDL-free portion of the substrate. Insets indicate the type of experiment. (H), (I), (J) Left: brightfield micrographs of control (top) and contracting edge (bottom) of the slice in intact CA3 (*n* = 8 control, *n* = 9 contracted), lesioned CA3 (*n* = 10 control, *n* = 10 contracted), and SUB (*n* = 9 control, *n* = 10 contracted), respectively. Right: schematics show slice placement on PDL-free surface in control (top) and contracted (bottom) groups. (K), (L), (M) Computed surface tension near the contracting edge of intact CA3, lesioned CA3, and SUB, respectively. Boxes indicate middle 50% range about the median. (two sample t-test: * p < 0.05, ** p < 0.01, *** p < 0.001).

**Fig. 4.**
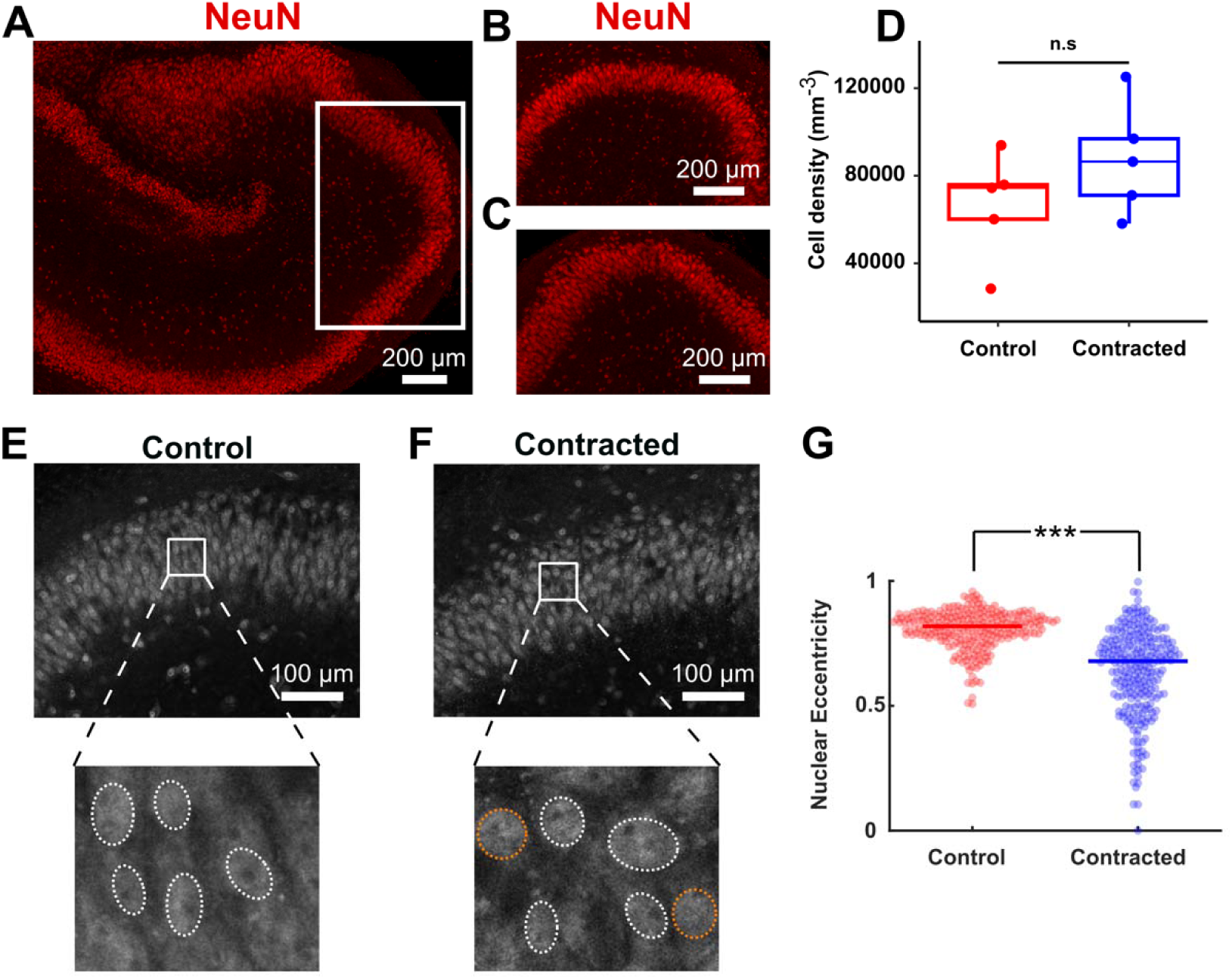
Cell density and morphological changes in contracted CA3 (intact). (A) Representative fluorescence micrograph of slice undergoing contraction in CA3, which was fixed on DIV 6 and stained with a marker for neuronal nuclei (NeuN). (B) Enlarged view of CA3 in a control slice (C) Enlarged view of contracted CA3, as indicated in (A) with a white box. (D) Cell density in control and contracted CA3 (*n* = 5 per group, two-sample t-test, n.s.: *p* = 0.222). (E), (F) Representative micrographs of NeuN staining in CA3 at higher magnification. In insets, nuclei with low and high eccentricity are indicated with white and orange dashed lines, respectively. (G) Nuclear eccentricity of neurons in control and contracted CA3 (*n* = 250 and 251 nuclei in control and contracte CA3, respectively, in *n* = 5 slices per group, Wilcoxon signed-rank test, ***p = 3.51 x 10^-37^, lines in the plot indicate medians of each distribution).

We cultured another group of slices on entirely PDL-free substrates. These slices exhibited a higher rate of contraction compared to control slices, with most PDL-free slices detaching from substrate by DIV 5 (Supplementary Fig. 2). We therefore did not use slices on entirely PDL-free substrates for further experiments.

We then compared the edges of slices cultured on substrates with patterned PDL coating to control slices. Regions of slices that underwent enhanced contraction (contracted group) had smoother edges compared to control group (Fig. 3H-J). Smoothness of tissue edge represents the degree of tissue surface tension due to cell compaction (Manning et al., 2010). Contracted regions showed an increase in surface tension at the contracting edge in all three configurations, compared to the control group (Fig. 3K-M).

We then examined the effect of contraction on neuronal morphology in CA3. Contraction altered the shape of neuronal nuclei, and a significant decrease in nuclear eccentricity was observed in the contracted slices (Fig 4 E-G).

### Neuronal activity in axially contracted CA3 (intact)

We studied the effect of accelerated contraction on different portions of the slice starting with intact CA3 undergoing contraction axially (Fig. 5A, B). Neuronal activity was monitored by expressing the Ca^2+^ indicator jRGECO1a (Dana et al., 2016) in neurons in the hippocampal slice culture, and measuring dynamic changes in fluorescence (ΔF/F). Activity was recorded on DIV 8, 13, 16, and 28, and seizure and non-seizure activity were analyzed separately. Area under the curve (AUC) and event frequency were calculated for non-seizure activity, revealing no significant differences between the contracted and control groups (Fig. 5 C-E). Slices with contracted CA3 tended to have lower seizure incidence compared to controls during DIVs 8, 13, and 16. However, on DIV 28, both groups showed a 100% seizure incidence (Fig. 5F). To compare seizure activity, we analyzed seizure amplitude, AUC, and total seizure duration (Fig. 5G). While all these seizure parameters exhibited a decrease in slices with contracted CA3 compared to controls, only the decrease in seizure AUC was statistically significant (Fig. 5 H, I, J). Overall trend was a decrease in seizure and non-seizure activity in axially contracted CA3.

**Fig. 5.**
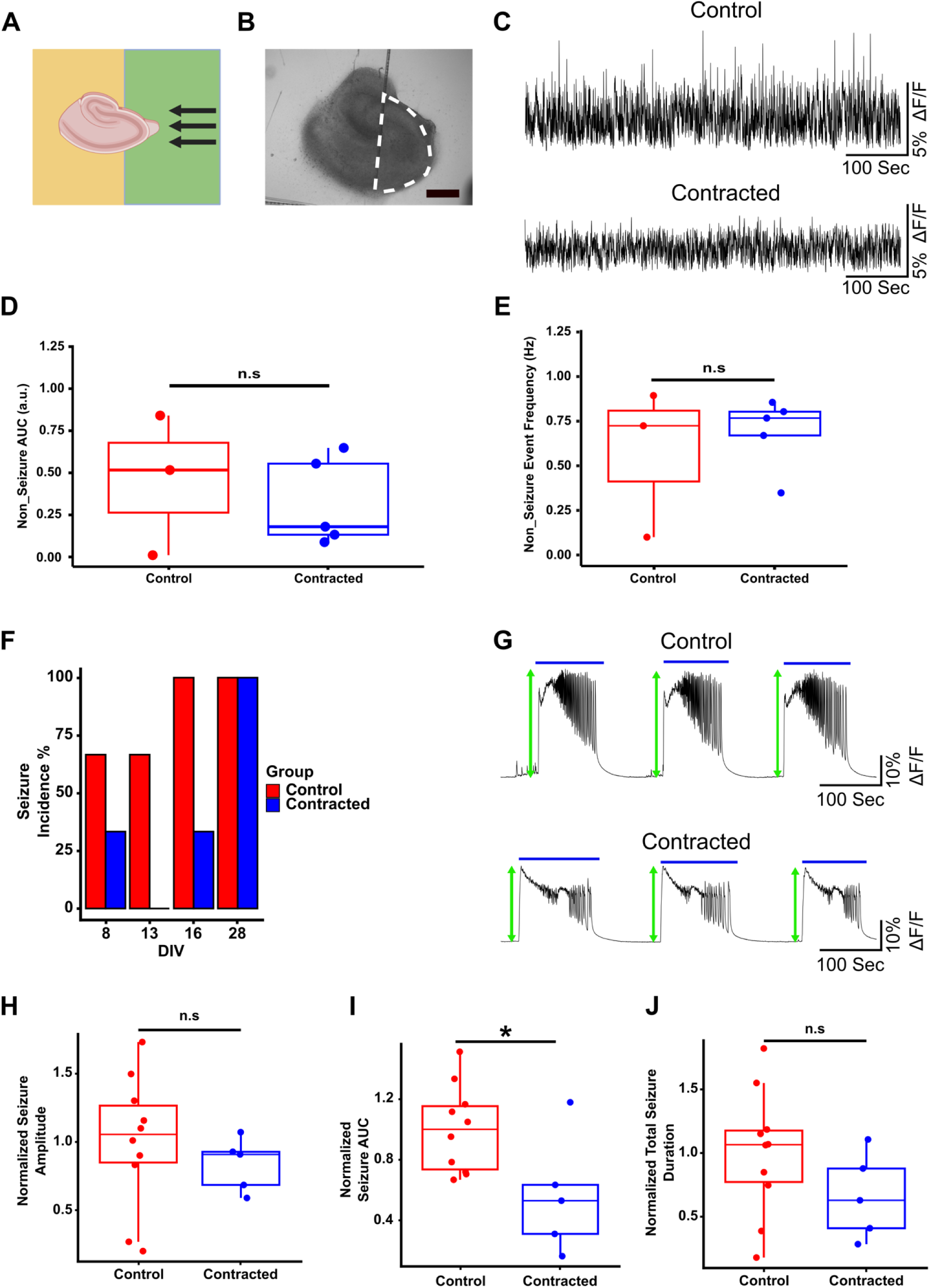
Neuronal activity in contracted CA3 (intact). (A) Schematic of slice configuration on the substrate with PDL pattern (green color corresponds to PDL-free region). (B) Bright-field image of the slice. White dashed line represents the recorded region of interest (ROI) (scale bar indicates 500 μm). (C) Representative examples of non-seizure neuronal activity from the same region in control and contracted slices. (D) Area under the curve (AUC) of non-seizure activity. (E) Event frequency of non-seizure activity. (F) Seizure incidence percentage in control and contracted groups on days of recording (*n* = 3 for both groups). (G) Representative examples of seizure activity from the same region in control and contracted slices. Blue solid lines indicate the detected seizures. Green arrows show seizure peaks (amplitude). (H) Normalized seizure amplitude in both groups. (I) Normalized seizure AUC in both groups (integration of detected seizures activity). (J) Normalized total seizure duration (summation of all detected seizure durations during each recording period) for both groups. (two-sample t-test: * p < 0.05, n.s: not significant).

### Neural activity in axially and transversely contracted CA3 (lesioned)

To study the effect of contraction in a different direction on neuronal activity, we removed the dentate gyrus (DG) and a portion of CA3, as shown in Fig. 6A. Slices with lesioned CA3 were positioned slices on a substrate with patterned PDL to apply contractile forces perpendicular to the apical-basal axis of neurons in the pyramidal layer. In this position, lesioned CA3 also experienced contractile forces along the apical-basal axis (Fig. 6A). Six regions of interest (ROI) were selected in CA3 for analysis of neuronal activity (Fig. 6B). Spontaneous activity was recorded on DIV 13, 18, and 20. The AUC of non-seizure activity was significantly higher in contracted slices compared to controls in some of the ROIs, while spike frequency trended higher, but was not significantly different (Fig. 6 C-E). Total seizure duration significantly decreased in the contracted group on DIV 13 but remained unaffected on other DIVs (Fig. 6H). Contracted slices displayed significantly higher seizure amplitudes compared to controls in all ROIs in CA3 (Fig. 6F, I), while seizure incidence and seizure AUC were not different between the two groups (Fig. 6G, J).

**Fig. 6.**
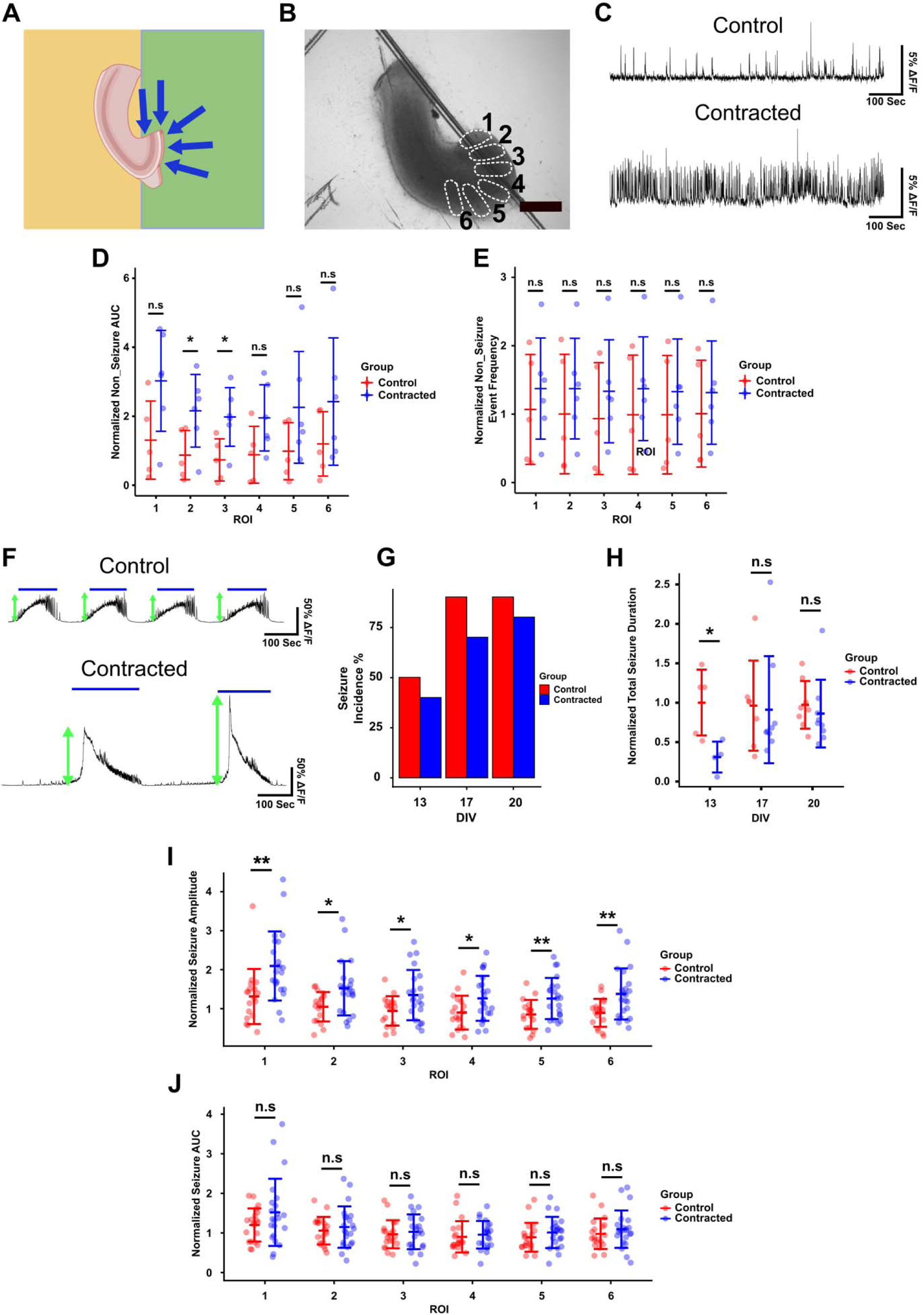
Neuronal activity in contracted CA3 (lesioned). (A) Slice configuration on the substrate with PDL pattern (green color corresponds to PDL-free region). (B) Regions of interest for recording activity in CA3 (scale bar indicates a distance of 500 μm). (C) Representative examples of non-seizure activity from ROI 1 in control and contracted slices. (D) Normalized area under the curve of non-seizure activity in the 6 ROIs. (E) Normalized event frequency of non-seizure activity in the 6 ROIs. (F) Representative examples of seizure activity from ROI 1 in control and contracted slices. Blue solid lines indicate the detected seizures. Green arrows show seizure peaks (amplitude). (G) Seizure incidence percentage in both groups and all recorded DIVs (n = 10 for both groups). (H) Total seizure duration in both groups for DIV of recording (summation of all detected seizure durations during each recording period). (I) Normalized seizure amplitude in both groups. (J) Normalized seizure AUC (integration of detected seizures activity) for both groups. (two sample t-test: ** p < 0.01, * p < 0.05, n.s: not significant).

### Neural activity in contracted subiculum and DG

We cultured hippocampal slices with subiculum and a portion of DG placed on PDL-free region of the substrate (Fig. 7A). We recorded neuronal activity in 6 ROIs in subiculum and 1 ROI in DG (Fig. 7B). There was no significant difference in AUC of non-seizure activity between contracted and control groups in any of the recorded ROIs in subiculum and DG (Fig. 7D, F). Non-seizure event frequency in the subiculum of contracted slices was lower but the difference was not statistically significant (Fig. 7E, Fig. S4). However, there was a significantly lower non-seizure event frequency in the contracted compared to control DG (Fig. 7G, Fig. S4). Seizure incidence was not significantly different between control cultures and cultures with contracted subiculum and DG (Fig. 7I). Seizure AUC was not affected by the contraction in subiculum (Fig. 7K), but significantly increased in DG compared to controls (Fig. 7L). In both DG and subiculum, seizure duration was comparable and unaffected by contraction (Fig. 7J). Seizure amplitude was significantly lower in most subiculum ROIs in contracted slices compared to controls (Fig. 7M), while no apparent difference was observed in seizure amplitude in DG (Fig. 7N). Overall, contraction resulted in small or no differences in neuronal activity in DG and subiculum.

**Fig. 7.**
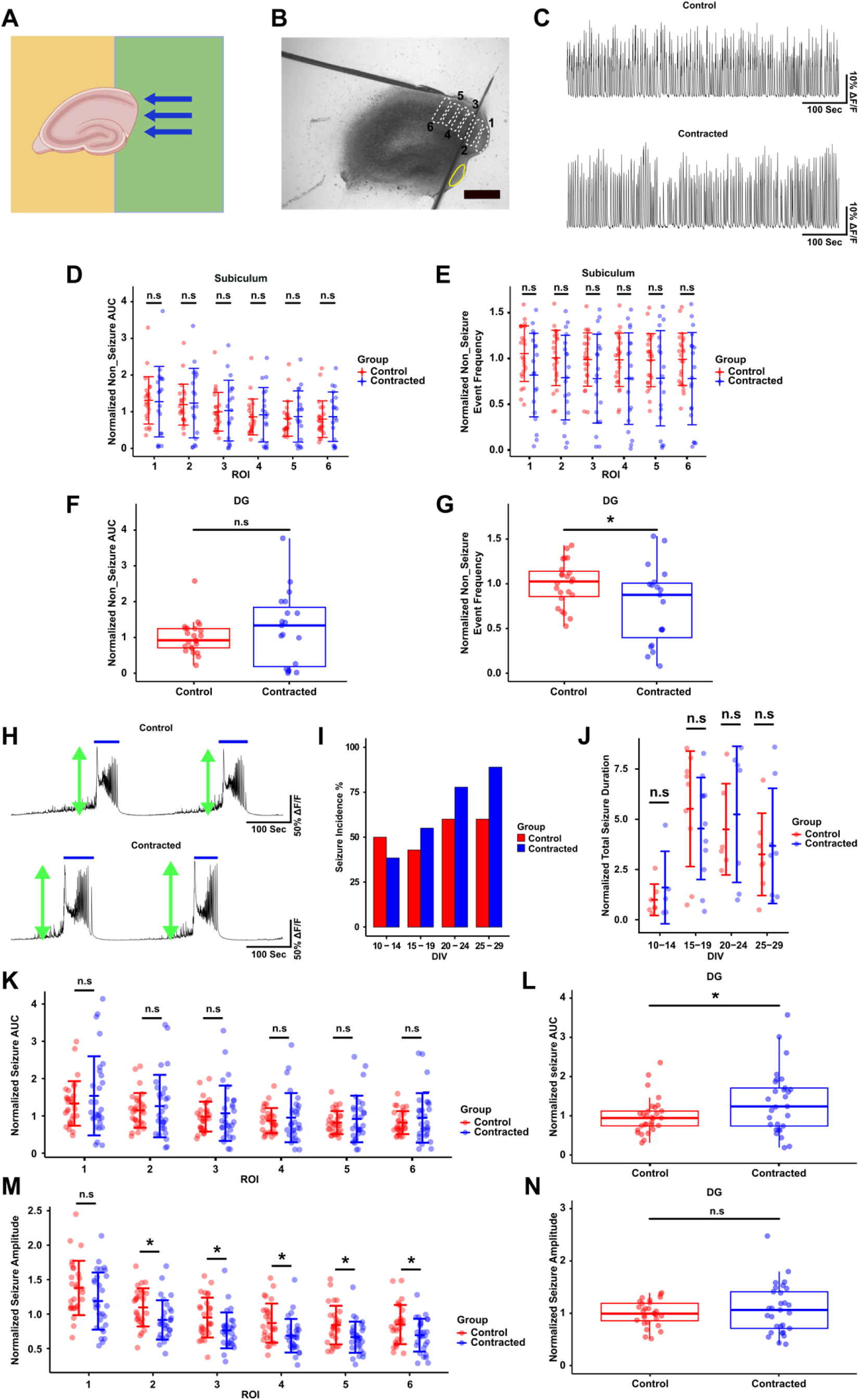
Neuronal activity in contracted subiculum and DG. (A) Configuration of the slice with subiculum and a portion of DG on the PDL-free region of the substrate (green). (B) Selected ROIs for neuronal activity recording in subiculum (white dashed line) and DG (yellow solid line). The scale bar indicates 500 μm. (C) Representative examples of non-seizure activity from ROI 1 of subiculum in control and contracted slices. (D) Normalized AUC of non-seizure activity in 6 ROIs in subiculum. (E) Normalized event frequency of non-seizure activity along 6 different ROIs in subiculum. (F) Normalized AUC of non-seizure activity in DG ROI. (G) Normalized event frequency of non-seizure in DG ROI. (H) Representative examples of seizure activity from subiculum ROI 1 in control and contracted slices. Blue solid lines indicate the detected seizures. Green arrows show seizure peaks (amplitude). (I) Seizure incidence percentage in both groups among all recorded DIVs (*n* = 14 controls, *n* = 13 contracted). (J) Total seizure duration in both groups on recorded DIVs (summation of all detected seizure durations during the recording period). (K) Normalized seizure AUC (integration of detected seizure activity) in both groups in subiculum ROIs. (L) Normalized seizure AUC in both groups in DG ROI. (M) Normalized seizure amplitude for both groups from subiculum ROIs. (N) Normalized seizure amplitude for both groups from DG ROI. (two-sample t-test: * p < 0.05, n.s: not significant).

## Discussion

Cells generate significant contractile forces through their cytoskeleton and actomyosin activity (Heer and Martin, 2017). These forces are transmitted through cell-cell junctions and cell-ECM adhesion points, resulting in tissue-level forces. Adult, healthy tissues exist in a state of mechanical homeostasis, where forces generated by various cells help to stabilize the tissue (Stamenović and L. Smith, 2020). In developing tissue, disbalance in cell-generated forces is responsible for processes such as the formation of Drosophila ventral furrow and the folding of the neural tube (Heer and Martin, 2017). In the developing brain, disbalanced mechanical forces drive folding of cortical tissue (Garcia et al., 2018). Disbalances of cell-generated forces during development are tightly orchestrated and guided by gene expression, and play an important role in tissue morphogenesis (Llinares-Benadero and Borrell, 2019). In contrast, disturbances in mechanical homeostasis occurring after development may have detrimental effects, and have been implicated in progression of atherosclerosis and tumor cell proliferation and invasion (Pillai and Franze, 2024; Stamenović and L. Smith, 2020). Atrophy of brain tissue after trauma may result in a disturbance of mechanical homeostasis in the surviving tissue, by perturbing the tissue-level balance of cell-generated forces. In an isolated slice of hippocampus, unbalanced tension generated by surviving cells can therefore be expected to produce forces pointing from slice periphery to slice center. Brain tissue is viscoelastic (Chatelin et al., 2010), suggesting that it will deform under sustained mechanical stress or tension. Under conditions of constant inward-pointing tension, the slice can be expected to undergo strong contraction – and this indeed occurred in our experiments (Fig. S2). Attachment of the slice to a substrate via poly-D-lysine introduced an artificial mechanical balance, and significantly reduced slice contraction (Fig. S2).

We then carried out experiments to confirm that slice contraction was an active process due to cell-generated forces. Tracking the movement of beads in the matrix surrounding the slice is a form of traction force microscopy (TFM). TFM is used to measure forces generated by individual cells or by engineered tissues (Obenaus et al., 2020). Forces generated by cells via cytoskeletal tension are transmitted to the surrounding matrix via integrin-mediated cell-matrix adhesions, and cause matrix deformation. Deformation can then be measured via bead displacement, with direction of bead movement indicating presence of tensile (movement of beads toward the tissue) or compressive (movement of beads away from the tissue) forces. Contractile tissues, such as engineered 3D epithelial tissues, were found to exert significant tensile forces, causing deformation of surrounding matrix, and movement of beads toward the portion of the tissue undergoing the largest contraction (Gjorevski and Nelson, 2012). Similar movement of beads toward contracting tissue can be observed in Fig. 2, indicating that Matrigel matrix is being deformed and pulled toward the slice. This may be caused by retracting slice edge pulling on the matrix via cell-matrix adhesions, as described above. An alternative view may be that bead movement indicates remodeling of Matrigel matrix by cells within the slice, either through secretion of ECM molecules, or through matrix degradation by matrix metalloproteases (Beroun et al., 2019; John et al., 2006; Krishnaswamy et al., 2019). However, production of new ECM could be expected to push beads away from the slice, while matrix degradation should result in release of beads into the culture medium. We have not observed these processes, and movement of beads toward the slice is consistent with the idea of cells within the slice generating tensile forces causing retraction of slice edges, in turn pulling the matrix toward the slice.

Our experiments with partial PDL coatings were designed to anchor one portion of the slice on PDL-coated region of the substrate, while allowing the portion of the slice above PDL-free region to contract freely. These experiments demonstrated that the contraction of hippocampal tissue was an active process shaped by the balance of mechanical forces. We observed that the portion of the slice that experienced the most contraction had well-defined, smooth edges. This effect was reminiscent of the effects of compaction observed in cellular aggregates (Yousafzai and Hammer, 2023). Cells in aggregate generate tensile forces via actomyosin contractily, and adhere to each other via cadherins on the cell surface. Combined effect of cell tension and adhesion results in the aggregate assuming a confirmation that minimizes its surface tension – a sphere with a smooth surface, with compacted individual cells (Manning et al., 2010). It appears that the portion of the hippocampal slice that experienced tensile forces unbalanced by PDL adhesion also underwent compaction, resulting in a smooth, curved tissue border (Fig. 3H-M). This suggests that mechanical interactions between hippocampal cells in ex vivo tissue may be governed by similar mechanisms as interactions between cells in aggregates. Hippocampal cells in regions of the slice that underwent retraction and compaction are also likely to have experienced significant mechanical force imbalances and morphological deformation. This could in turn lead to hyperexcitability through mechanotransduction pathways.

Mechanotransduction, or conversion of mechanical stimuli into cellular signals, can occur via multiple pathways in neurons. These pathways include initiation of cell signaling cascades via force-sensing cytoskeletal proteins, opening of mechanosensitive ion channels, and alteration of transcription due to mechanically-induced deformation of nuclear envelope (Pillai and Franze, 2024). One or more of these mechanisms may be involved in sensation of unbalanced mechanical forces by neurons in the retracted, compacted region of hippocampal slice. We have directly observed deformation of the nuclear envelope of CA3 neurons due to contraction, suggesting that this mechanism of mechanotransduction may be responsible for observed excitability changes. Furthermore, deformation of neuronal soma, dendrites, and axons by tissue-level retraction and compaction may also alter neuronal excitability. We found that excitability may increase or decrease depending on the orientation of the contractile forces to basal-apical axis of CA3 neurons. This suggests that mechanotransduction through dendrites may play a significant role in the effects we have observed. A limitation of our study is that we have not identified the specific mechanotransduction mechanism that links cellular forces to changes in excitatability.

Spontaneous seizure-like activity appears in organotypic hippocampal cultures after one to two weeks post-isolation. This reflects the time course of epileptogenesis in this model system. When compared to the time course of mechanically-driven events described in this work, it appears that spontaneous seizures follow slice contraction after a time lag. Tissue contraction may therefore play a role in epileptogenesis. Organotypic hippocampal cultures are a complex, heterogeneous system composed of multiple cell types that include excitatory and inhibitory neurons, astrocytes, microglia, and other cells. Cells respond to the injury of dissection and isolation from the rest of the brain by axonal sprouting in neurons (Berdichevsky et al., 2013), and inflammatory activation of astrocytes and microglia (Chong et al., 2018; Magalhães et al., 2018). These processes: network reorganization due to axon sprouting and formation of excessive connectivity, and inflammation-driven effects of glia on neuronal function, may be significant contributors to epileptogenesis in organotypic hippocampal cultures. Mechanically-driven tissue contraction, and resulting changes in excitability, occur concurrently with sprouting and inflammation in this model system. Experiments where a portion of the slice was placed on non-adhesive, PDL-free surface, enabled us to compare slices with a contracting region to control slices. Both experimental groups can be expected to experience inflammation and sprouting, but only slices with a contracting region experienced additional cell-generated mechanical forces. This allowed us to isolate mechanically-driven changes in excitability from those driven by sprouting and inflammation. We observed increase in excitability during both seizure and non-seizure activity due to cell-generated forces directed orthogonally to apical-basal axis of CA3 neurons, but a decrease in excitability during seizure-like activity when forces were directed parallel to apical-basal axis of CA3 neurons or orthogonal to apical-basis axis of subicular neurons. These changes occurred in a system where neuronal excitability is also driven by network reorganization and inflammation. It is possible that some of the region-specific differences we observed could be due to maximization of excitability by these mechanisms of epileptogenesis. In other words, excitability in subicular neurons may already be near maximum value, making it difficult to observe effects of cell-generated forces on excitability during non-seizure activity, for example. Further study in models with reduced intrinsic excitability may shed light on this question.

## Conclusions

We have used cultures of hippocampal slices as a model of disbalance in mechanical forces that may occur in the brain after trauma. We found that cell generated forces in this model result in contraction and compaction of hippocampal tissue. We also found that changes in tissue geometry in turn result in excitability changes in a region-specific manner. Results of experiments reported in this study suggest that imbalanced cell-generated forces contribute to development of epilepsy, and represent a novel mechanism of epileptogenesis after trauma.

## Methods

### Slice preparation

Organotypic slices were prepared from the hippocampi of Sprague-Dawley rat pups that were dissected on post-natal days 7-8. The McIlwain tissue chopper (Mickle Laboratory Eng. Co., Surrey, United Kingdom) was used to chop the hippocampi into slices with 350 μm width. The slices were cultured on a 35 mm culture-treated petri dish (Thermo Fisher) partially or completely coated with 0.5 mg/ml poly-D-lysine (PDL) solution (Sigma Aldrich, Cat # P0899) in 0.1M Borate buffer (pH 8.5). The culture medium for the slices consisted of 97.5% Neurobasal-A media (Gibco, Cat# 10888-022), 2% B-27 supplement (Gibco, Cat# 17504-044), 0.25% 200 mM GlutaMAX (Gibco, Cat#35050-061), and 0.3% gentamicin (Gibco, Cat# 15710-064). This medium was added to the slices and replaced every 3-4 days with fresh medium. The cultures were incubated at 37°C in environment with 5% CO_2_ on a rocker. All animal use protocols were approved by the Institution Animal Care and Use Committee (IACUC) at Lehigh University and were conducted in accordance with the United States Public Health Service Policy on Humane Care and Use of Laboratory Animals.

### Slice cultures embedded in bead-containing Matrigel

Fluorescent latex beads with 1 μm diameter (Sigma, L4655) were diluted in Neurobasal-A medium at a ratio of 1:2000. Then, 2 μL of this solution was mixed with 150 μL of Matrigel (Corning, Cat # 354234) in a microtube on ice. After removing the culture medium from slices on DIV 1, 5 μL of Matrigel and beads mixture was added on top of the slices and kept at room temperature without medium for 10 minutes to let the Matrigel gel. The culture medium was then added. To track the movement of the beads, fluorescent and bright field images were captured daily using an inverted fluorescence microscope (IX73, Olympus) equipped with a CCD camera (Thorlabs) and aligned using the registration plugin (Linear stack alignment with SIFT) in Fiji (ImageJ). ImageJ ROI tools were used to measure the beads’ movement distance and angle.

### PDL patterns and contraction measurement

Polydimethylsiloxane (PDMS) films were prepared by mixing liquid PDMS and the curing agent (Sylgard 184 by Electron Microscopy Sciences, Cat # 24236-10) at a 10:1 ratio and baking overnight at 60° C in an oven (Thermo Fisher). Square polydimethylsiloxane (PDMS) structures with an embedded square well were cut from the PDMS film. These PDMS structures were then placed onto the culture substate (tissue-culture treated Petri dishes). PDL solution was then poured into the well to coat the exposed dish surface and incubated overnight at 37^°^ C. The dish was then washed, and the borders of the PDL-coated area were marked on its surface before detaching the PDMS. The PDMS was carefully removed, and organotypic slices were cultured with regions intended to have no surface adhesion placed over the PDL-free portions of the substrate. Control slices were cultured on dishes entirely coated with PDL. The cultured slices were imaged daily using an inverted microscope (CKX41, Olympus) and the area of contracted ROI over PDL-free portion of the substrate in contracted slices and same ROI in control slices were measured in ImageJ. Slice contraction in the selected ROI was calculated according to the Eq. 1; where A1 is the initial ROI area and A6 is the ROI area on DIV 6.

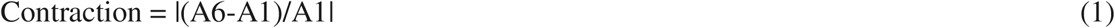

### Surface tension analysis at the edge of slices

The method is shown in Supplementary Fig. 3. In ImageJ, pixel values were extracted along a line perpendicular to the contracting edge of the slice. These values were then plotted versus distance along the line. A sigmoid curve was fitted to the data using a 4-parameter logistic model in MATLAB curve fitter. R-square values and visual inspection was used for goodness-of-fit assessment. R^2^ > 0.9 was considered acceptable. The surface tension was calculated as the inverse of the distance between the maximum and minimum values of the fitted sigmoid curves.

### Immunohistochemistry, cell counting, and nuclear eccentricity analysis

Cultured slices were fixed at DIV 7 in 4% paraformaldehyde (Electron Microscopy Sciences, Cat # 15710) for 2 hours and subsequently washed and permeabilized in 0.3 % Triton X-100 (Sigma-Aldrich, Cat # T8787) in Dulbecco’s phosphate-buffered saline (Sigma-Aldrich, Cat # D8662) on a shaker for 2 hours. Following this, the slices were blocked with 10% goat serum (Gibco) overnight at 4° C. Anti-NeuN antibody conjugated to Alexa Fluor 555 (Millipore Sigma, Cat # MAB377A5) at a 1:100 dilution ratio was then added to the samples, which were kept in 4° C for 72 hours on a shaker. The stained slices were mounted on a microscope slide using one drop of Fluoro-Gel (Electron Microscopy Sciences, Cat # 17985-10). Samples were imaged using a confocal microscope (Zeiss LSM 510 META, Germany) with a 20X objective to acquire z stacks at 0.96 μm intervals. Confocal images were analyzed in Fiji (ImageJ), and cell counting of NeuN-positive cells in CA3 was performed using Fiji cell counter plugin in a rectangular region of fixed size for all slices.

Nuclei of NeuN-positive cells were identified, and their long and short axes were measured in Fiji (ImageJ). Long axis was defined as the axis bisecting the nucleus with direction closest to the normal to the pyramidal layer, while short axis was defined as axis closest to the tangential to the pyramidal layer. Lengths of long (d_long_) and short (d_short_) axes were used to calculate eccentricity for each nucleus using Eq. 2.

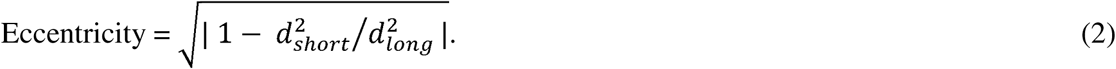

### Calcium imaging

JRGECO1a, a genetically encoded [Ca^2+^] indicator, was expressed under the Synapsin promoter by adding pAAV.Syn.NES-jRGECO1a.WPRE.SV40 (titer ≥ 5×10 ^9^ vg/mL) to the culture medium on DIV 1. pAAV.Syn.NES-jRGECO1a.WPRE.SV40 was a gift from Douglas Kim & GENIE Project (Addgene plasmid # 100854) (Dana et al., 2016).

Changes in jRGECO1a fluorescence, corresponding to neural activity, were observed using an inverted fluorescence microscope (IX73, Olympus) while the culture dish was placed in a mini-incubator (Bioscience Tools) supplied with humidified blood gas and maintained at 37° C. Fluorescence changes were recorded by a CCD camera (Thorlabs) at 5 frames/sec rate for 20 minutes. The mean fluorescence intensity was calculated for each frame within the regions of interest (ROIs) in ImageJ. Change in fluorescence was calculated using Eq. 3:

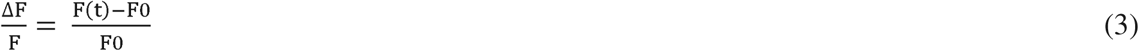

where F(t) represents the raw mean fluorescence in the ROI, and F0 is the baseline calculated in MATLAB employing the asymmetric least square mean smoothing method (Eilers and Boelens, 2005).

### Neural activity analysis

Recorded neural activity was classified into seizure and non-seizure categories for analysis. Seizures were manually detected as prolonged paroxysmal events that occurred across all regions of interest with at least 10% ΔF/F and lasted at least 10 seconds. The seizure incidence was determined by calculating the percentage of cultures within each group that exhibited at least one seizure during the 20-minute recording period. Area under the curve for the entire signal was calculated and divided by the recording duration. Non-seizure portions of the recording (that did not contain seizures or oscillations) were filtered by high-pass Butterworth filter using the filtfilt function in MATLAB Signal Processing Toolbox. The filter was 4th-order, with cutoff frequencies ranging from 0.05 Hz to 0.25 Hz. Non-seizure area under the curve (AUC) was then calculated by summing the area under the curve of filtered and thresholded signal, and dividing it by recording duration. Threshold was determined as 3*standard deviation of ΔF/F in silent regions. Seizure AUC was determined by summing the area under the curve for all identified seizures in a recording, and dividing it by the total recording period. Total seizure duration was calculated by summing the duration of all detected seizures in a recording divided by the total recording period. We determined the seizure amplitude by taking the average peak of all detected seizures in each recording. Non-seizure event frequency was calculated in MATLAB by detecting ΔF/F peaks at various thresholds above the noise, and selecting the maximum value as the corresponding frequency.

## Statistical analysis

The distribution of all data sets was tested for normality using Shapiro-Wilk or K-S test. Two-sample t-test was used for unpaired data sets and paired t-test was used for a paired data set. Chi-squared test with Pearson correction was performed to compare the distribution of bead movement angles with a uniform distribution. A significance level of 5% was considered for determining statistical significance. The Wilcoxon signed-rank test was used to assess statistical significance in nuclear eccentricity. The type and results of statistical analysis are indicated in figure legends.

## Data availability

All data supporting the findings of this study are available and will be provided upon request to the corresponding author.

## Acknowledgments

All schematic diagrams used in the figures were created with Biorender.com.

## Author contributions: CRediT

LD: Methodology, Investigation, Data curation, Formal analysis, Visualization, Writing – original draft, STL: Funding acquisition, Supervision, Writing – review and editing, YB: Conceptualization, Methodology, Formal analysis, Project administration, Funding acquisition, Supervision, Validation, Writing – original draft.

## Supplementary information

Supplementary Figures S1 – S4.

