## Supplementary Figures for "Cell-generated mechanical forces play a role in epileptogenesis after injury"

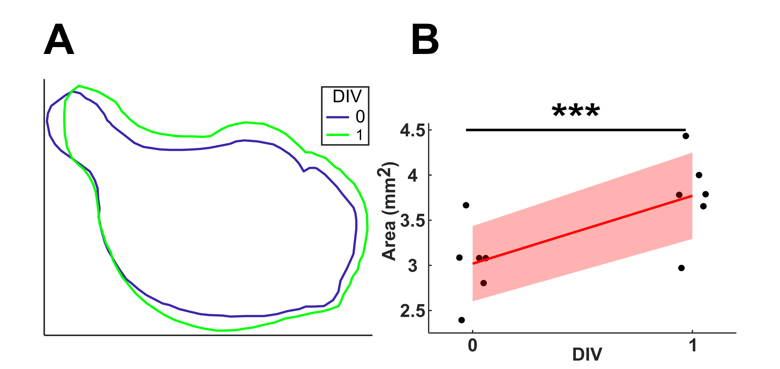

**Fig. S1 Natural contraction of organotypic slices during first day of culture.** (A) Boundary of slice on DIV 0 (day of dissection) and DIV 1. (B) Calculated area of cultured slices (n = 6) from DIV 0 to DIV 1. Paired t-test was used to measure statistical significance (*** p < 0.001).

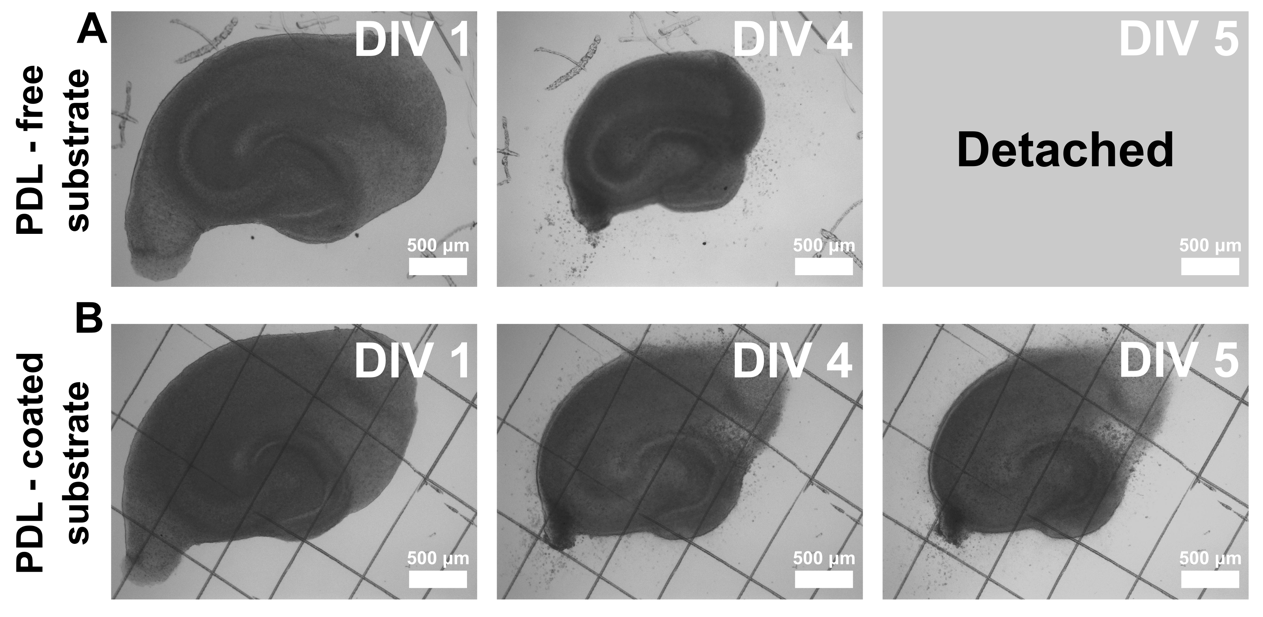

**Fig. S2 Comparison of hippocampal slice cultures on PDL-free and fully PDL-coated substrates.** (A) from left to right: bright field images of an organotypic hippocampal slice cultures on PDL-free substrate at DIV1, DIV4, and day of detachment (DIV5) (B) from left to right: bright field images of an organotypic hippocampal slice on entirely PDL coated substrate at DIV1, DIV2, DIV5.

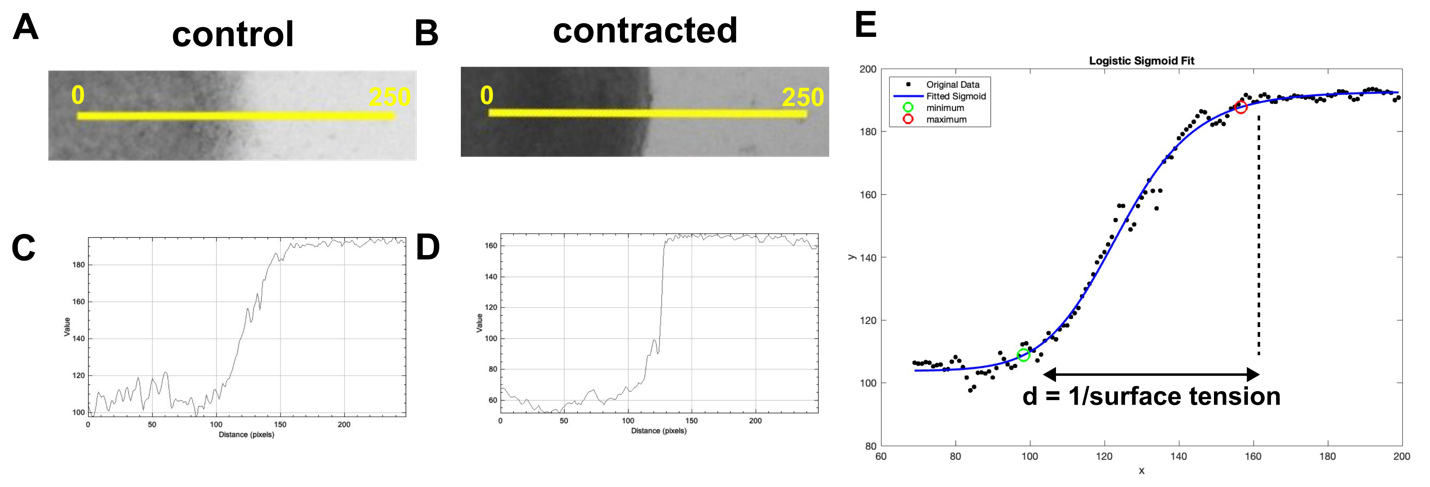

**Fig. S3 Surface tension analysis method at the edge of slice.** (A) Edge of subiculum in control slice. Yellow solid line was drawn perpendicular to the slice edge and was used for pixel value extraction at slice edge. (B) Edge of subiculum in contracted slice. (C) Pixel values plotted versus the distance along the yellow solid line in (A). (D) Pixel values plotted versus the distance along the yellow solid line in (B). (E) Pixel values versus distance (dots) with the corresponding sigmoid-fitted curve (blue line). The minimum and maximum values are indicated with green and red circles, respectively. Surface tension at the slice edge was calculated as the inverse of the distance between the minimum and maximum of the fitted curve.

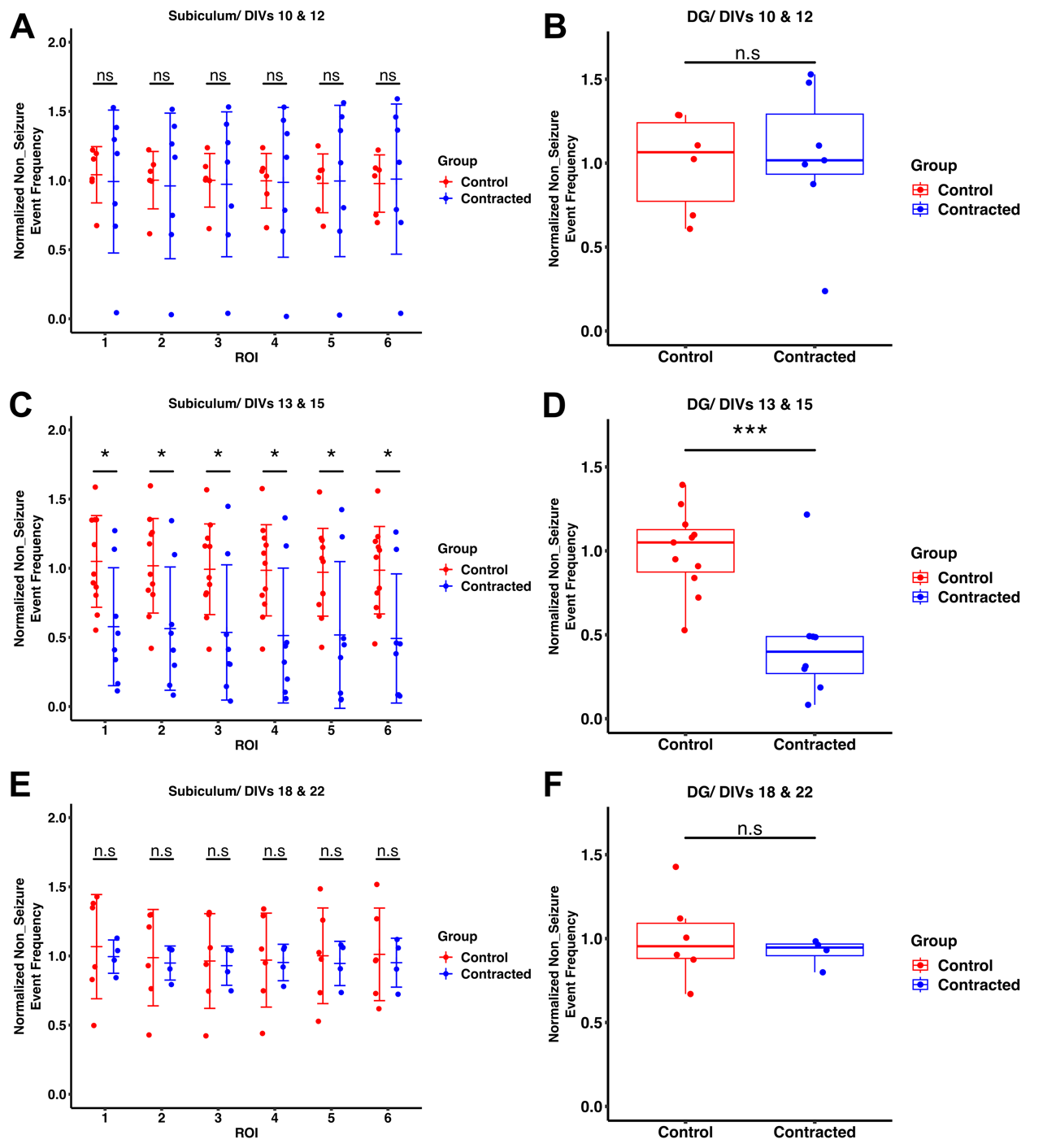

**Fig. S4: Non-seizure activity event frequency in the subiculum and DG over time.**
(A), (C), (E) Normalized event frequency across six ROIs in the subiculum at DIV10/12, DIV13/15, and DIV18/22. (B), (D, (F) Normalized event frequency in the DG ROI at the same time points. (two-sample t-test: *** p < 0.001, * p < 0.05, n.s: not significant).
